# Specific Detection of Endogenous S129 Phosphorylated α-Synuclein in Tissue Using Proximity Ligation Assay

**DOI:** 10.1101/2021.09.28.461511

**Authors:** Ryan Arlinghaus, Michiyo Iba, Eliezer Masliah, Mark R Cookson, Natalie Landeck

## Abstract

**BACKGROUND:** Synucleinopathies are a group of neurodegenerative disorders that are pathologically characterized by the accumulation of protein aggregates called Lewy Bodies. Lewy bodies are primarily composed of α-synuclein (asyn) protein, which is phosphorylated at serine 129 (pS129) when aggregated. Currently available commercial antibodies used to stain for pS129 asyn can cross react with other proteins, thus making it difficult to specifically detect endogenous pS129 asyn and to interpret pS129 asyn staining.

**OBJECTIVE:** To develop a staining procedure that detects pS129 asyn with high specificity and low background.

**METHODS:** We use the fluorescent and brightfield in situ Proximity Ligation Assay (PLA) to specifically detect pS129 asyn in cell culture, mouse and human brain sections.

**RESULTS:** The pS129 asyn PLA specifically stained for endogenous, soluble pS129 asyn in cell culture, mouse brain sections, and human brain tissue without significant cross-reactivity or background signal.

**CONCLUSIONS:** This PLA method can be used to specifically detect pS129 asyn in order to further explore and understand its role in health and disease.

## Introduction

Dementia with Lewy Bodies, Parkinson’s disease, and Multiple system Atrophy are a group of neurodegenerative diseases termed Synucleinopathies. Synucleinopathies are pathologically characterized by the presence of α-synuclein (asyn)-positive protein inclusions in the brain called Lewy Bodies (LBs) and lewy neurites (Spillantini et al., 1997; Spillantini & Goedert, 2000). Prior work has established that the asyn found in LBs is post-translationally modified through nitration, truncation, ubiquitination, SUMOylation, and phosphorylation (Fujiwara et al., 2002; Giasson, Duda, et al., 2000; W. Li et al., 2005; Shahpasandzadeh et al., 2014; Tofaris et al., 2003). Of these post-translational modifications, the phosphorylation of asyn at serine 129 (pS129) is the most extensively studied. Under normal cellular conditions pS129 asyn comprises less than 1% of total asyn, whereas about 90% of aggregated asyn is phosphorylated at this residue (Fujiwara et al., 2002; Landeck et al., 2016). Therefore, pS129 asyn has been used as a marker of asyn aggregation and pathology in synucleinopathies and has additionally been explored as a biomarker for disease progression (Chen & Feany, 2005; Foulds et al., 2011; Fujiwara et al., 2002; Gómez-Tortosa et al., 2000; Ortuño-Lizarán et al., 2018; Stewart et al., 2015; Wang et al., 2012). However, several aspects of pS129 biology are unclear, including whether phosphorylation of asyn directly influences asyn toxicity, whether it increases or decreases asyn aggregation propensity *in vivo*, or whether it is simply a byproduct of other pathological processes (see review: (Oueslati, 2016)).

Given the strong ties between pS129 asyn and the pathology of synucleinopathies, it is critical to have tools that specifically detect pS129 asyn. Previously developed monoclonal antibodies raised against the pS129 asyn epitope preferentially detect pS129 asyn over non-phosphorylated asyn. However, it has been challenging to limit the cross-reactivity of pS129 asyn monoclonal antibodies with other proteins that have similar phosphorylated epitopes, including neurofilament light chain (Delic et al., 2018; Rutherford et al., 2016; Sacino et al., 2014; Uchihara & Giasson, 2016). The significant cross-reactivity of these monoclonal antibodies makes it a challenge to specifically detect and interpret the results from pS129 asyn staining.

Proximity Ligation Assay (PLA) is a staining method that creates a robust signal when two primary antibodies are within an estimated range of <40nm (Söderberg et al., 2006). This range depends on the size of the primary and secondary antibodies, as well as the length of the oligonucleotides attached to the secondary antibodies. When the two conjugated oligonucleotides are close together, they can be ligated and subsequently amplified to create a fluorescent or a 3,3’-Diaminobenzidine (DAB) signal. PLA can therefore be used to demonstrate the close proximity of two proteins within a cell (Fredriksson et al., 2002; Söderberg et al., 2006). Additionally, PLA can be used to improve detection of protein modifications by using one antibody targeted to the protein and one targeted to the modification. This approach limits the cross-reactivity and background signal of primary antibodies (Hegazy et al., 2020; Lindskog et al., 2020; Zieba et al., 2018)). In the asyn field, PLA has also been used to stain oligomeric asyn by using a single asyn antibody, and also to detect the association of asyn and α-tubulin in mice and human striatum (Amadeo et al., 2021; Roberts et al., 2019). Here, we describe a novel PLA system to stain for pS129 asyn using a pS129 asyn antibody MJF-R13 (abcam) and a total-asyn antibody syn-1 (BD Biosciences). We show that our novel PLA system specifically detects pS129 asyn in mouse brain sections and primary neuron cultures with lower non-specific staining compared to conventional primary antibodies. Additionally, we illustrate how this PLA method is sensitive enough to detect endogenous pS129 asyn in cultured primary neurons, mouse and human brain sections.

## Results

### Assessment of pS129 α-synuclein antibody specificity using conventional immunostainings

To evaluate whether commercially available pS129 asyn antibodies are monospecific, we performed immunocytochemistry (ICC) on mouse primary cortical neurons. We used the pS129 asyn monoclonal antibodies MJF-R13 (abcam), pSyn#64 (WAKO), and 81A (abcam) (Fig 1A-C,E-G) (Waxman & Giasson, 2008). A total-asyn antibody, syn-1 (BD Biosciences), was used as a reference (Fig 1D,H) and experiments were performed with both asyn knock-out (SNCA KO; Fig 1A-D) and wild-type (WT; Fig 1E-H) cultures.

**Figure 1:**
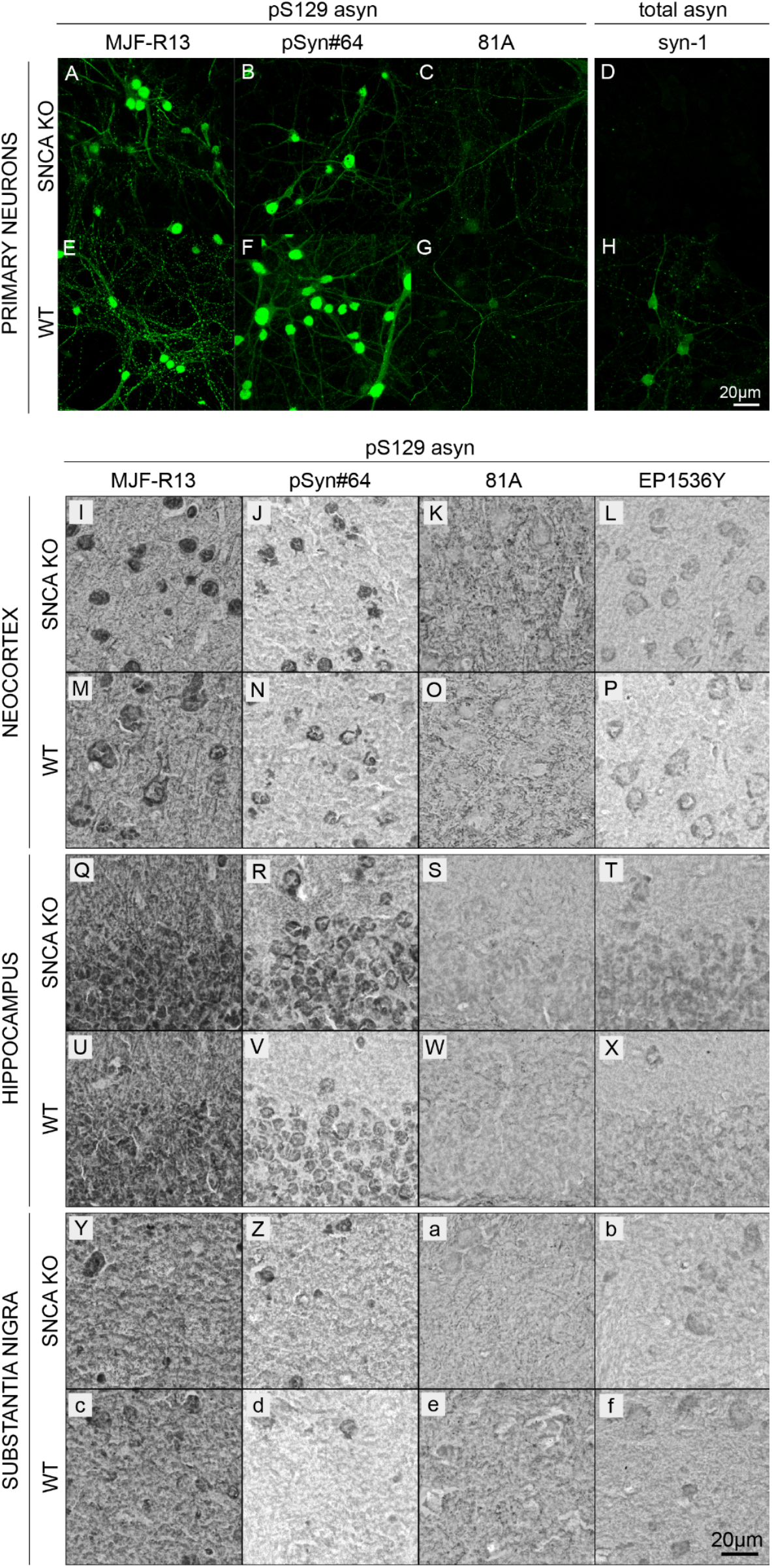
Cross-reactivity of pS129 α-synuclein monoclonal antibodies in cell culture and mouse brain sections. (A-H) Immunocytochemistry staining (ICC) of SNCA KO (**A-D**) and WT (**E-H**) mouse primary cortical neurons using rabbit MJF-R13 (**A,E**), mouse pSyn#64 (**B,F**), and mouse 81A (**C,G**). Total-asyn antibody, mouse syn-1 (**D,H**), was used as a control. Alexa-488 stained images were taken at 63x. (I-f) Immunohistochemistry staining (IHC) of SNCA KO (**I-L,Q-T,Y-b**) and WT (**M-P,U-X,c-f**) 30μm thick coronal mouse brain sections with rabbit MJF-R13 (**I,M,Q,U,Y,c**), mouse pSyn#64 (**J,N,R,V,Z,d**), mouse 81A (**K,O,S,W,a,e**), and rabbit EP1635Y (**L,P,T,X,b,f**). DAB-stained images were taken at 63x.

MJF-R13, pSyn#64, and 81A antibodies showed extensive staining in both the WT and SNCA KO cultures, demonstrating cross-reactivity of pS129 asyn monoclonal antibodies with other antigens. In contrast, the syn-1 antibody produced a staining in the WT culture only, illustrating that this antibody is monospecific in this application.

Next, we evaluated the specificity of the pS129 asyn monoclonal antibodies MJF-R13, pSyn#64, 81A, and EP1536Y (abcam) using immunohistochemistry (IHC) (Fig 1I-f). IHC was performed on paraformaldehyde (PFA) fixed coronal SNCA KO (Fig 1I-L,Q-T,Y-b) or WT (Fig 1M-P,U-X, c-f) 30μm mouse brain sections containing the cortex, hippocampus, and substantia nigra. In all three brain regions, there were no differences in staining patterns between WT and SNCA KO samples. For antibodies MJF-R13, pSyn#64 and EP1536P somatic staining was visible along with background staining throughout all brain regions. The 81A antibody did not show cell body staining and had more signal intensity in the cortex and substantia nigra than other areas.

In summary, none of the commercially available antibodies that we used here specifically detect endogenous levels of pS129 asyn in ICC or IHC.

### Staining of pS129 α-synuclein in primary cortical neurons and mouse brain sections using the Proximity Ligation Assay

We next aimed to develop an assay to detect pS129 asyn that would work equally well in rodent and human tissues. Many asyn antibodies that are commercially available recognize the C-terminus of asyn, and typically these antibodies do not recognize both human and rodent asyn with the same affinity due to sequence differences in this region (Croisier et al., 2006; Vaikath et al., 2019). The syn211 and LB509 clones are human specific monoclonal antibodies that bind very close to the pS129 asyn site (Giasson, Jakes, et al., 2000; Jakes et al., 1999) and were not considered due to potential competition with pS129 antibodies. The 4B12 and syn204 clones recognize epitopes more distant to S129, but were also eliminated as they do not recognize rodent asyn (Giasson, Jakes, et al., 2000; Landeck et al., 2016). We therefore considered C-20 (Santa Cruz) and syn-1 (BD Biosciences) asyn antibodies that can recognize human and mouse total asyn with similar affinity (Landeck et al., 2016; Perrin et al., 2003). However, C-20 is no longer commercially available, which led us to choose the mouse monoclonal syn-1. Syn-1 is produced in mice and recognises amino acids 91-99 of asyn (Fig 2A) (Perrin et al., 2003). To achieve a specific PLA signal, we required a monoclonal pS129 asyn antibody with high affinity and therefore chose the MJF-R13 (R13) clone which has no preference for either rodent or human asyn (Landeck et al., 2016). An overview of the PLA protocol is shown in Fig 2A-D. To evaluate if our PLA specifically stains for pS129 asyn in mouse primary cortical neuron cultures, we performed the fluorescent version of the PLA (Fluorescent-PLA) in SNCA KO (Fig 2E), WT (Fig 2F), and WT primary cultures overexpressing mouse asyn (masyn) or human asyn (hasyn) mediated by adeno-associated viral vectors (AAV) (Fig 2G and H, respectively). pS129 asyn monoclonal antibody R13 and total asyn monoclonal antibody syn-1 were used for the fluorescent-PLA at a concentration of 1:1000 and 1:2000, respectively. Our Fluorescent-PLA produced a signal in the WT culture that was specific to asyn as demonstrated by the minimal signal detected in the SNCA KO cultures. Signal intensity was further increased in the masyn and hasyn overexpression WT cultures (Fig 2G,H). These data demonstrate that our Fluorescent-PLA is capable of specifically staining pS129 asyn in primary neuron cultures and that it can be used to detect endogenous levels of pS129 asyn.

**Figure 2:**
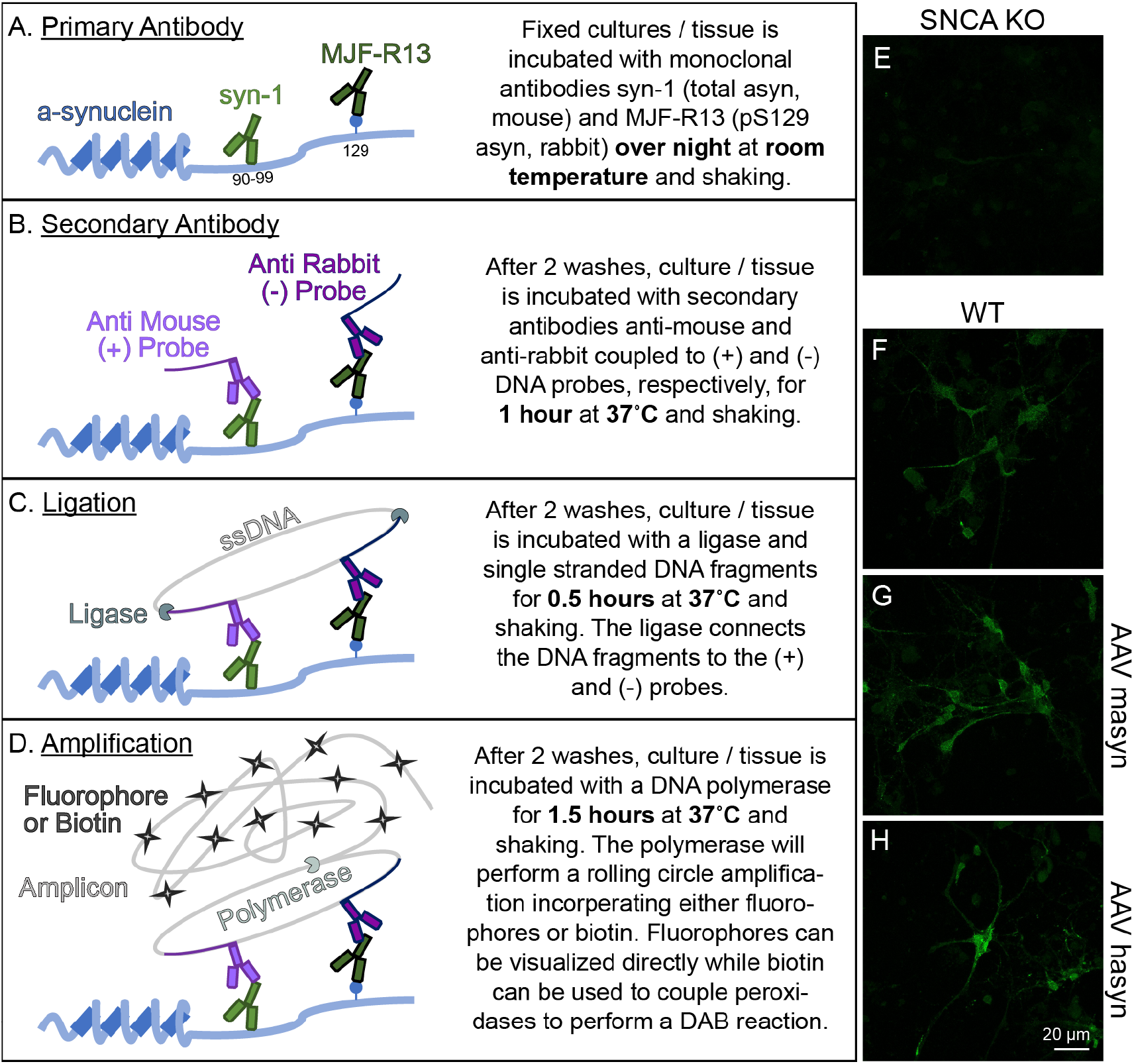
Fluorescent-PLA specifically detects pS129 α-synuclein in primary cortical neuron cultures. (A-D) Schematic illustration of the principle experimental steps in the PLA protocol. (E-H) The fluorescent-PLA specifically stains for pS129 asyn in WT (**F**) but not in SNCA KO (E) primary cortical neuron cultures. AAV overexpression of mouse asyn (AAV masyn) (G) and human asyn (AAV hasyn) (H) in WT cultures were used to further exemplify the utility of the Fluorescent-PLA.

We then evaluated the brightfield version of the PLA (BF-PLA) utilizing DAB to stain for pS129 asyn on PFA fixed brain tissue (Fig 3). pS129 asyn antibody R13 (1:1,000) and total-asyn syn-1 (1:2,000) were used as the primary antibodies. The BF-PLA was performed with both primary antibodies (Fig 3A-H) or with only one of the primary antibodies, either syn-1 (Fig 3I-L) or R13 (Fig 3M-P) alone. We used WT (Fig 3A-D, I-P) and SNCA KO (Fig 3E-H) coronal mouse brain sections (30μm) to evaluate the specificity and sensitivity of our pS129 asyn PLA in the neocortex, striatum, hippocampus, and substantia nigra. When both primary antibodies were used, the BF-PLA produced a robust, punctate signal in the WT neocortex, striatum, hippocampus, and substantia nigra (Fig 3A-H). The punctate staining is expected as PLA staining relies on rolling circle amplification to produce amplicons that limit diffusion of chromogen from the stained area. Dramatically less signal was produced in the SNCA KO tissue in any of the brain regions (Fig 3E-H). The syn-1 antibody used alone demonstrated very little nonspecific staining in all brain areas (Fig 3I-L) while the sections stained with just R13 alone showed small numbers of punctae, mainly in the hippocampus (Fig 3M-P). These findings illustrate that our BF-PLA can be utilized to specifically stain for pS129 asyn in mouse brain sections and that this BF-PLA shows higher specificity than traditional DAB protocols using pS129 asyn monoclonal antibodies.

**Figure 3:**
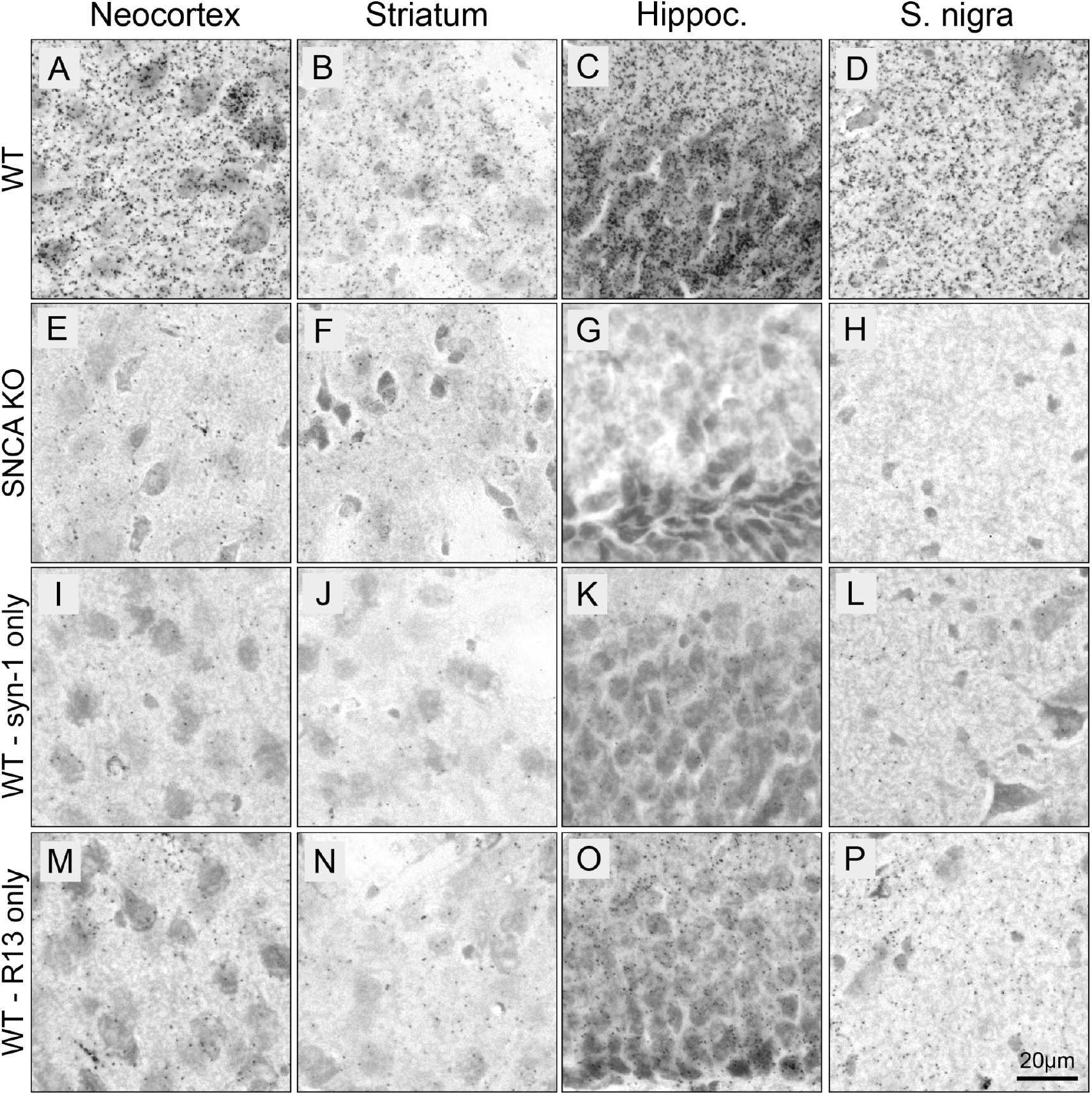
BF-PLA specifically detects pS129 α-synuclein in mouse brain sections. **(A-P)** BF-PLA staining of pS129 asyn was performed on coronal mouse brain sections (30μm). R13 (1:1,000) and syn-1 (1:2,000) monoclonal antibodies were used together (**A-H**). The BF-PLA was also performed using either syn-1 (**I-L**) or R13 (**M-P**) alone as controls. WT (**A-D,I-P**) or SNCA KO sections (**E-H**) containing neocortex (**A,E,I,M**), striatum (**B,F,J,N**), hippocampus (**C,G,K,O**), and substantia nigra (**D,H,L,P**) were used.

Next, we evaluated the ability of the BF-PLA to stain for endogenous pS129 asyn in additional major brain areas in the WT mouse brain. (Fig 4). A punctate staining was noted throughout the stained mouse brain sections. In the prefrontal cortex (PFC), similar to the neocortex, the staining was not only seen throughout (Fig 4A,C) but was particularly strong in layer IV, likely due to staining of neuronal cell bodies (Fig 4E). As expected, the pS129 asyn staining was homogenous throughout the striatum and there was no obvious staining of cell bodies (Fig 4F-G). The hippocampus was mainly stained in the CA3 region with cell bodies being labeled intensely (Fig 4H-I). Additional somal staining was visible especially in the paraventricular nucleus of the thalamus (Fig 4J-K), as well as in the amygdala (Fig 4L-M). Even though asyn is well known to be highly expressed in dopaminergic neurons in the substantia nigra and total staining can be visible in their cell bodies (Supplementary Fig 1W,X), we did not see an increased staining of pS129 asyn in the reticulata or pars compacta subregions (Fig 4N-O). Additional intense staining was seen in the granular layer of the cerebellum where cell bodies were visible with varying strength of staining (Fig 4P-Q).

**Figure 4:**
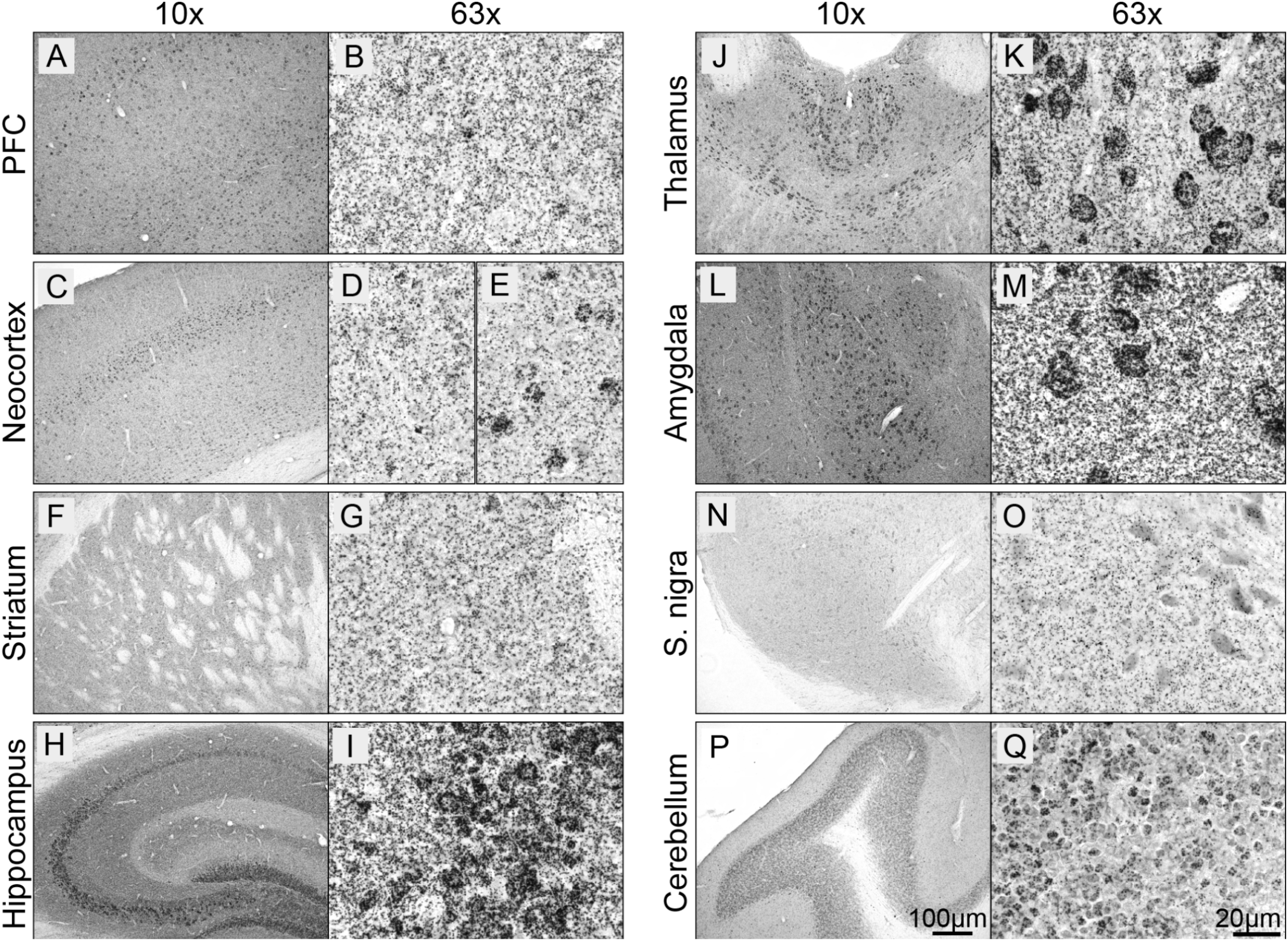
Endogenous pS129 α-synuclein staining in the mouse brain. (A-Q) WT coronal mouse brain sections (30μm) were stained using the BF-PLA technique. Examples of prefrontal cortex (PFC; **A,B**), neocortext (**C-E**), striatum (**F,G**), hippocampus (**H,I**), thalamus (**J,K**), amygdala (**L,M**), substantia nigra (S. nigra; **N,O**) and cerebellum (**P,Q**) are shown in 10x (**A,C,F,H,J,L,N,P**) and 63x (**B,D-E,G,I,K,M,O,Q**) magnification.

These data show that we are able to detect endogenous pS129 asyn in primary mouse neurons and in mouse brain sections using fluorescent-PLA and BF-PLA, respectively. To our knowledge, this is the first time endogenous pS129 asyn has been detected in mouse brain tissue without a nonspecific signal. We also demonstrate that pS129 asyn is found throughout the mouse brain and that in specific brain areas there is strong pS129 asyn staining in cell bodies.

### pS129 α-synuclein staining in α-synuclein overexpression animal models and in human brain tissue using BF-PLA

Since pS129 asyn is often used as a marker of asyn pathology, we also wanted to validate our BF-PLA staining of pS129 asyn in relevant animal models (Fig 5). First, we performed BF-PLA staining on mouse brain sections from WT mice overexpressing rodent or human asyn (Fig 5A-L). WT mice received unilateral injections into the substantia nigra with AAV expressing either GFP (Fig 5B,F,J), mouse asyn (masyn; Fig 5C,G,K), or human asyn (hasyn; Fig 5D,H,L). Four weeks post-injection, we used BF-PLA to stain for pS129 asyn in the substantia nigra and striatum. Uninjected brain sections of WT mice were used in parallel (Fig 5A,E,I). Both the AAV-masyn and AAV-hasyn showed increased staining intensity in the substantia nigra compared to the uninjected and AAV-GFP brain sections (Fig 5G,H). The additional pS129 asyn signal was mainly found in cell bodies and not in the projections of neurons located in the surrounding tissue or in the striatum (Fig 5I-L). No obvious differences were seen between the uninjected and control AAV-GFP stainings (Fig 5E,F), suggesting that the increase in staining intensity is not due to the AAV injections but rather due to an increase in asyn expression. We then investigated the BF-PLA staining of pS129 asyn in WT mice that received injections of masyn preformed fibrils (PFF). PFFs were injected into the striatum and animals were euthanized 1 month post injection. In line with previous studies, extensive pS129 asyn staining was seen in the striatum, neocortex, prefrontal cortex, and the amygdala of paraffin embedded sections (Fig 5M-T). While endogenous, soluble pS129 asyn signal throughout the brain was visible, we were also able to detect accumulation of pS129 asyn in cell bodies in all four regions, with particularly high expression in the amygdala (Fig 5Q-T, black arrows).

**Figure 5:**
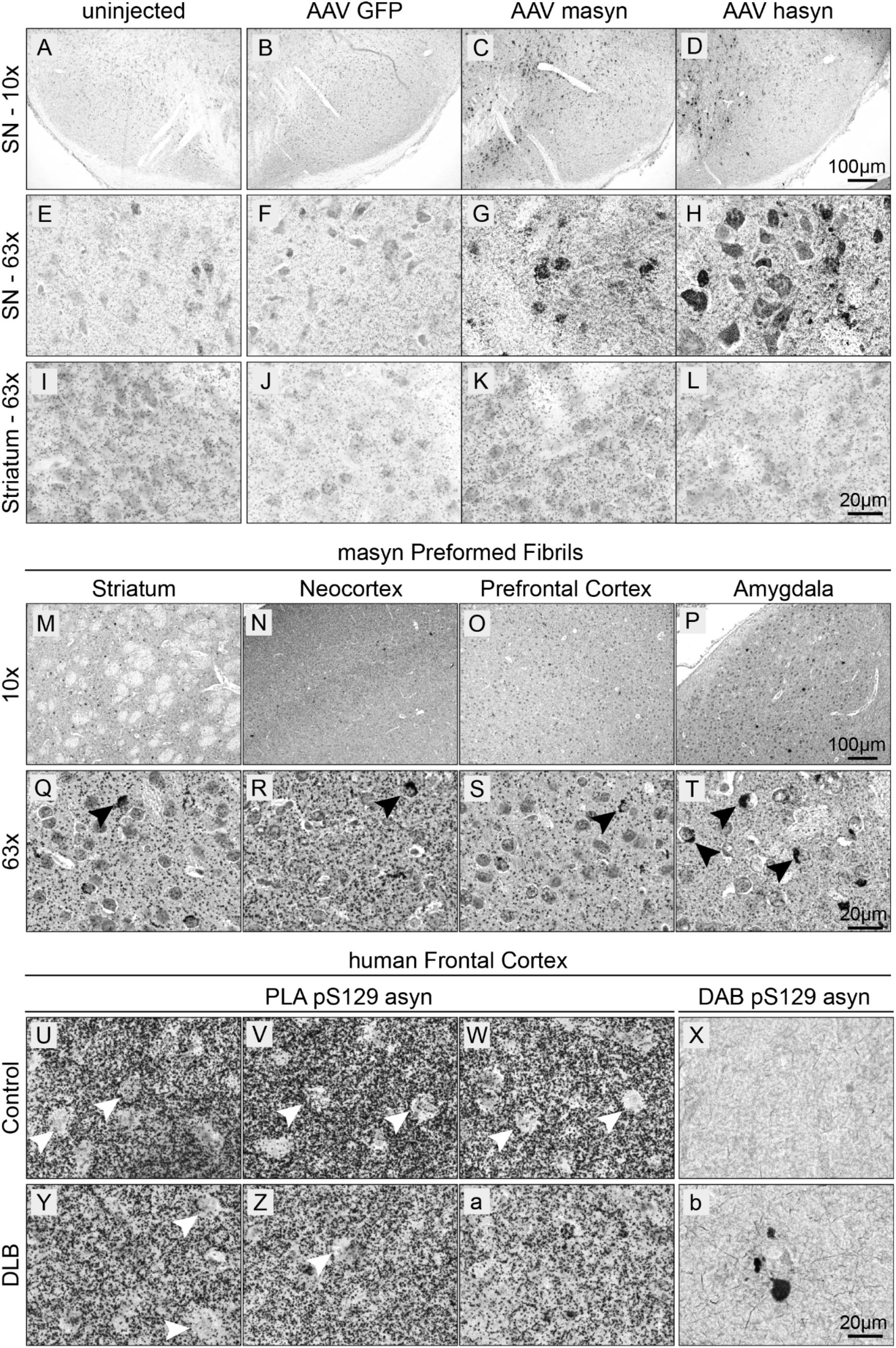
BF-PLA used on sections from animal models and human frontal cortex. (**A-L**) Mice were injected with AAV expressing either GFP (**B,F,J**), mouse asyn (masyn) (**C,G,K**) or human asyn (hasyn) (**D,H,L**). Four weeks after injection, the brains were fixed, cut into 30μm thick sections, and stained using the BF-PLA. Uninjected sides were used as a baseline control (**A,E,I**). Images were taken either with 10x (A-D) and 63x (**E-L**) magnification. (**M-T**) Mice were injected with asyn PFF into the striatum. One month after injection, the mice brains were fixed, embedded in paraffin, cut into 6μm thick sections, and stained using the BF-PLA. Images of striatum (**M,Q**), neocortex (**N,R**), prefrontal cortex (**O,S**) and amygdala (**P,T**) were taken at 10x (**M-P**) and 63x (**Q-T**) magnification. Accumulation of pS129 asyn staining in cell bodies indicated by black arrows (**Q-T**). (**U-b**) Human frontal cortical sections from a healthy control (**U-X**) and a DLB patient (**Y-b**) were acquired from ADRC and stained using the BF-PLA protocol. Images were taken at 63x magnification. The monoclonal pS129 asyn antibody, 81A, was used singly to confirm the presence and absence of LBs in the DLB patient tissue and healthy control tissue, respectively (**X,b**).

Next, we evaluated the use of BF-PLA to detect soluble pS129 asyn in human brain tissue, includings aggregated pS129 asyn in LBs. Brain tissue from a neurologically unimpaired control and a patient with Dementia with Lewy Bodies (DLB) was acquired from the Alzheimer Disease Research Center (ADRC) at the University of California, SanDiego (UCSD). In both the control and DLB human brain tissue, BF-PLA showed extensive pS129 asyn staining in frontal cortical areas (Fig 5U-W and Y-a, respectively). Interestingly, minimal staining was observed in cell bodies suggesting that pS129 asyn is primarily located in projections in the human frontal cortex (Fig 5U-W,Y-Z, white arrows). Although there was less staining of projections in the DLB patient tissue (Fig 5Y-a), we could not confirm this observation when we stained three additional cases for each group (Supplementary Fig 1A-L). Additionally, our BF-PLA did not readily detect LBs in the DLB human brain tissues (Fig 5Y-a, Supplementary Fig 1B,F,J,D,H,L). To confirm the presence and absence of LBs in the DLB patient samples and healthy control samples, we used the 81A antibody for DAB staining of pS129 asyn. pS129 asyn aggregates were found in the DLB patient tissue while none were detected in the healthy control (Fig 5X,b). We performed additional staining using the total human asyn specific 4B12 combined with R13 to address whether steric hindrance of the syn-1 antibody prevented it from detecting the tightly packed aggregated pS129 asyn. Of note, the combination of 4B12 and R13 has been successfully used in a previously described assay for pS129 asyn (Cariulo et al., 2019) and has proven to be human and asyn specific in our hands (Supplementary Fig 1O,P). As with the syn-1/R13 BF-PLA, we detected soluble pS129 asyn readily, but we did not detect any LB inclusions (Supplementary Fig 1M,N). Taken together, our BF-PLA can be successfully used in animal models to detect accumulated and soluble endogenous pS129 asyn. Additionally, the BF-PLA is capable of staining endogenous pS129 asyn in human brain tissue. However, the BF-PLA assay was not able to detect densely packed asyn in LBs in the tissue of a human patient with DLB.

## Discussion

Here, we present a novel tool to detect pS129 asyn with high specificity. We describe a PLA protocol that specifically detects endogenous pS129 asyn in cell culture, in mouse brain sections with validation using knockout tissue and in human brain tissue, including brain tissue from DLB patients. We also show that this PLA protocol can be used to detect increased levels of pS129 asyn in cell culture and mouse models overexpressing asyn, as well as PFF injected mice. Due to its specificity in detecting pS129 asyn, as well as its utility in disease models, this PLA may have utility in answering some of the major questions still surrounding pS129 asyn biology and its pathological role in synucleinopathies.

While using the Sigma-Aldrich Duolink^®^ PLA kit, we noticed that staining variations were seen between different kits and batches. We believe this may be in part due to the low-temperature activated Ligase and Polymerase enzymes used during the protocol, which might make them more vulnerable at room temperature. Therefore, special care needs to be taken when using these enzymes. Vials should always be kept in cryo containers and only as short as possible at room temperature. Additional staining differences were seen depending on the experimental setup. When staining 30μm thick brain sections with the BF-PLA, only a thin upper layer (≈ 5μm) of the section was successfully stained, especially if the section was already mounted before staining and not free floating. Sections were mostly stained already mounted to glass slides to reduce the amount of reagents needed. Better penetration and staining was seen for 6μm thick paraffin embedded sections. These penetration issues could come from the antibodies or enzymes not being able to successfully penetrate deep into the sections. Further optimization could be done by using varying amounts of detergent, different fixative, or antigen retrieval to improve staining thickness. While the protocol for our pS129 asyn PLA is more time intensive compared to traditional ICC or IHC and uses two antibodies produced in mouse and rabbit, it is still possible to co-stain other antigens on the same tissue. In brain sections, immuno-alkaline phosphatases and fast blue can be used as long as the primary antibody is not produced in mice or rabbits. Likewise, fluorescent PLA stainings only require one fluorophore and thus other antigens can be stained if the antibodies are not hosted in mouse or rabbit.

We show here, for the first time, the distribution of pS129 asyn throughout the mouse brain and its cellular localization *in vivo* without major cross-reactivity of the monoclonal antibodies. Interestingly, in the mouse brain, pS129 asyn can be located as a general signal in most parts of the brain but also as more dense staining specifically in some, but not all, cell bodies. This could either suggest a different function of pS129 asyn in distinct neurons or that phosphorylation of asyn is more prominent in cells that highly express the relevant kinases. It is worth noting that the localization of these somal stainings of pS129 asyn in the various subregions do not necessarily overlap with the localization of pathology seen in synucleinopathies (Beach et al., 2009; Jellinger, 2004; Saito et al., 2003). In the human frontal cortex, we saw a slightly different pattern. While there was again a general staining in every part of the frontal cortex section, we did not see a localization of pS129 asyn in most of the soma for the cases used here. Quantification of pS129 asyn levels using immunohistochemistry have in general been tricky as these techniques can be easily influenced by many factors such as fixation, post mortem delay and localization on slides. The PLA method has even more components such as ligation and amplification. Therefore, we recommend to use this assay only for assessment of protein localization.

One drawback of the R13 / syn-1 PLA is the lack of staining of pathological aggregated pS129 asyn in human brain tissue. As the assay does stain accumulated pS129 asyn in the PFF animal model, we hypothesize that this may be due to the chronically aggregated, dense LBs having a different structure or different post-translational modifications when compared to the short-term aggregated asyn in animal models. The syn-1 antibody binds near the 90th amino acid on asyn, which is part of the motif involved in building the β-sheet structure in asyn aggregates (Y. Li et al., 2018) and includes a ubiquitination site at K96 (Schmid et al., 2013). Both these modifications may thus prevent syn-1 from binding to asyn that is aggregated in LBs but is still able to detect endogenous and soluble human pS129 asyn. To rectify this discrepancy, we performed the BF-PLA using 4B12 for total asyn combined with R13 on human DLB tissue. We again detected soluble but not aggregated pS129 asyn. This could either be because of a sterical hindrance of the 4B12 and syn-1 antibodies binding to LB or due to R13 detecting only a certain set of post-translational modifications besides pS129 that are only found on soluble but not aggregated asyn. Several studies have found that besides phosphorylation on S129, asyn has various other modifications such as ubiquitination, truncation, nitration, SUMOylation, and phosphorylations, for example at Y125 (El Turk et al., 2018; Giasson, Duda, et al., 2000; W. Li et al., 2005; Shahpasandzadeh et al., 2014; Tofaris et al., 2003). Any of these modifications may influence the binding of antibodies, especially antibodies that are sensitive to post translational modifications, like R13. More work needs to be done in the future to better characterize the exact epitopes of pS129 asyn antibodies.

Overall, our pS129 asyn PLA with the R13 and syn-1 monoclonal antibodies shows higher specificity than stainings with pS129 asyn antibodies alone, and is also capable of detecting the low endogenous levels of pS129 asyn in mouse and human tissue.

## Conclusion

The PLA described here addresses the issue of cross-reactivity of pS129 asyn monoclonal antibodies by establishing a specific signal for pS129 asyn with minimal background signal. We show how this PLA protocol can be used to specifically detect endogenous pS129 asyn in cell culture, mouse brain sections, and human brain tissue. Furthermore, we demonstrate its use in cell culture and mouse models to detect increased levels of pS129 asyn. While soluble pS129 asyn was readily detected in human brain tissue, LBs were not stained by our PLA protocol. Due to its diverse utility in specifically detecting pS129 asyn *in vitro* and *in vivo*, our PLA can be utilized to answer major questions still surrounding pS129 asyn biology and its role in the pathology of synucleinopathies.

## Methods

### Primary Cultures

Postnatal day 0 WT or SNCA KO cortices were dissected in ice cold HBSS. The cortices were then incubated in papain for 30 min at 37 °C. After incubation, cortices were washed twice with BME (Basal Medium Eagle + 0.45% glucose + 0.5 mM glutamate + Pen/Strep + N2 + B27) with centrifugations at 1000 rpm for 3 min. Tissue was triturated in BME containing 50 μg/ml of DNAse and then washed twice in BME with centrifugations at 1000 rpm for 6 min. Cells were diluted with BME + 5% Fetal Bovine Serum (FBS) to a concentration of 1.0×10^6 cells/ml. 1.0×10^6 cells were added to each well in a 24-well plate with poly-D-lysine - Laminin coated coverslips. The 24-well plate was then placed in a 37 °C and 5% CO_2_ incubator. After 24 hours, 80 % of BME + 5 % FBS was removed and replaced with fresh BME containing 2.5 μM Cytosine Arabinoside. DIV 6 primary neurons were treated with human asyn or mouse asyn adeno-associated virus (AAV) 7×10^13 vg/ml for 4 hours. Cultures were not kept past Day In Vitro (DIV) 9. Primary neurons were fixed by addition of 4 % PFA for 30 min, washed and stored in 1x PBS at 4 °C.

### Animals

Strains used are C57Bl/6J (WT; Jax) and C57Bl/6J 129S6-Snca^tm1nbm^ (SNCA KO;(Cabin et al., 2002)). Mice were housed in a 12-hour light/dark cycle with *ad libitum* access to food and water. All animal work was performed in accordance with the Institutional Animal Care and Use Committee (ACUC) of the National Institute on Aging (NIA/NIH).

### Stereotaxic injection - viral vectors

Animals were anesthetized by 4% isoflurane. After placing the animal into a stereotaxic frame (Kopf), 1 μL of rAAV6 vector solution was unilaterally injected into the substantia nigra (SN) using the following coordinates: anteroposterior (AP) −3.2 mm, mediolateral (ML) −1.5 mm and dorsoventral (DV) −4.3 mm from bregma. The tooth bar was adjusted to −0.6 mm. Injection was performed using a pulled glass capillary attached onto a 5 μl Hamilton syringe with a blunt 22s gauge needle. After delivery of the viral vector using a pulsed injection of 0.1 μl every 15 s, the capillary was held in place for 5 min, retracted 0.1 mm and after 1 min it was slowly withdrawn from the brain. Ketoprofin was administered s.c. as analgesic treatment for 3 days post op.

### Mouse brain sections

Animals were killed when < 1 year old or 4 weeks after viral vector injection by an overdose of ketamine and perfused via the ascending aorta first with 10 ml of 0.9% NaCl followed by 50 ml of ice-cold 4% paraformaldehyde (PFA in 0.1 mM phosphate buffer, pH 7.4) for 5 min. Brains were removed and post-fixed in 4% PFA for 24 h and then transferred into 30% sucrose for cryoprotection. The brains were then cut into 30 μm thick coronal sections and stored in an antifreeze solution (0.5 mM phosphate buffer, 30% glycerol, 30% ethylene glycol) at −20 °C.

### Stereotaxic injection and sections - PFF

5μg of asyn PFF (2μg/μl, 2.5μl) was injected to induce α-synucleinopathy (Luk et al., 2012) into the striatum (anteroposterior (AP) 0.2 mm, mediolateral (ML) 2 mm and dorsoventral (DV) −3.2 mm from bregma) of wild type mouse, after deeply anesthetized with isoflurane and immobilized in a stereotaxic frame under aseptic conditions. The animal was observed during and after the surgery and Ketoprofin was administered for three days including the surgery day. Mouse was harvested at 1 month post injection period, perfused with PBS and the brain was fixed with 70% Ethanol in PBS. Then brains were cut into 2-mm thick slices and embedded in paraffin after overnight paraffinizing. The paraffin block was sectioned with 6μm thickness. Before the PLA staining, the slide was deparaffinized with xylene and alcohol series and pretreated with a citrate buffer for antigen retrieval.

### Human brain tissue

Human frontal cortex and hippocampus samples from DLB cases and neurologically unimpaired controls were obtained from the Alzheimer Disease Research Center (ADRC) at the University of California, SanDiego (UCSD). Brain tissue was fixed with formalin post mortem and later cut into 40 μm thick free floating sections using the vitratome.

### Immunocytochemistry

Neurons were blocked in 5% FBS in PBS for 30 min, and incubated with primary antibodies, diluted 1:500 in 1% FBS/PBS overnight. Following three washes in PBS, the cells were incubated for 1 h with a secondary antibody diluted 1:500 (Alexa Fluor 488-conjugated; ThermoFisher) in 1% FBS/PBS solution. After three PBS washes, the coverslips were mounted and the cells were analyzed by confocal microscopy (Zeiss LSM 880 microscopy).

### Immunohistochemistry

Sections were washed with PBS and incubated for 30 min in blocking buffer (10% Normal Donkey Serum (NDS), 1% BSA, 0.3% Triton in PBS). Afterwards, primary antibodies were used at 1:500 and incubated overnight in 1% NDS, 1% BSA, 0.3% Triton in PBS. Next day, sections were washed three times with PBS and incubated with 1:500 Alexa Fluor 488-conjugated secondary antibody for 1h at room temperature. After three washes with PBS, sections were mounted on glass slides, coverslipped using Prolong Gold Antifade mounting media (Invitrogen) and imaged using a Zeiss LSM 880 confocal microscope equipped with Plan-Apochromat 63X/1.4 numerical aperture oil-objective (Carl Zeiss AG).

### Antibodies

**Table 1:** Antibodies used to demonstrate cross-reactivity of pS129 asyn monoclonal antibodies and those used for the PLA to specifically detect pS129 asvn are listed below with the Drovider information and antibodv specifics

| Antibody | Epitope | Company | Catalog # | Host Species | Clonality | Specific detection of: |  |  |
| --- | --- | --- | --- | --- | --- | --- | --- | --- |
|  |  |  |  |  |  | Human asyn | Rodent asyn | pS129 asyn |
| Syn1 | 91-99 | BD Biosciences | AB_398107 | Mouse | Monoclonal, IgG1 | Yes | Yes | No |
| 4B12 | 103-108 | Covance Inc. | SIG-39730-500 | Mouse | Monoclonal, IgG1 | Yes | No | No |
| MJF-R13 |  | Abcam | ab168381 | Rabbit | Monoclonal, IgG | Yes | Yes | Yes |
| EP1536Y |  | Abcam | ab51253 | Rabbit | Monoclonal, IgG | Yes | Yes | Yes |
| 81A |  | Abcam | ab184674 | Mouse | Monoclonal, IgG2a | Yes | Yes | Yes |
| pSyn#64 |  | WAKO | 014-20281 | Mouse | Monoclonal, IgG1 | Yes | Yes | Yes |

### Proximity Ligation Assay (PLA)^®^ Sigma Aldrich – Millipore

#### Reagents

Duolink^®^ In Situ PLA^®^ Probe Anti-Rabbit PLUS Affinity purified Donkey anti-Rabbit IgG (H+L) (DUO92002), Duolink^®^ In Situ PLA^®^ Probe Anti-Mouse MINUS Affinity purified Donkey anti-Mouse IgG (H+L) (DUO92004), Duolink^®^ In Situ Detection Reagents Green (DUO92014), Duolink^®^ In Situ Detection Reagents Brightfield (DUO92012), Duolink^®^ In Situ Wash Buffers, Fluorescence (DUO82049)

#### Protocol

was generally followed as described by the manufacturer. Briefly, for BF-PLA, sections were incubated in H_2_O_2_ solution for 30 min at room temperature free floating and shaking followed by 2 washing steps with Wash Buffer A. Always ensure that Wash Buffer A is at room temperature before use. All sections were mounted, dried and incubated in Duolink^®^ Blocking buffer for 1 hour. All incubation steps were performed in a humidity chamber at 37°C with gentle shaking (50 rpm to ensure an even staining) unless stated otherwise. Afterwards, samples were incubated overnight at room temperature with primary antibodies anti-pS129 (ab209421 MJF-R13, Abcam) and total anti-asyn (42/α-Synuclein syn-1, BD Biosciences) 1:1000 and 1:2000, respectively, diluted in Duolink^®^ Antibody Diluent. The next day, samples were washed 2x for 5 minutes with gentle shaking using Wash Buffer A. Secondary antibodies Duolink^®^ In Situ PLA^®^ Probe Donkey Anti-Mouse IgG (H+L) MINUS and Duolink^®^ In Situ PLA^®^ Probe Donkey Anti-Rabbit IgG (H+L) PLUS were diluted 1:5 in Duolink^®^ Antibody Diluent and incubated for 1 hour. After 2x 5 minute washes with gentle shaking using Wash Buffer A, sections were incubated in 1x Duolink^®^ Ligation Buffer and Ligase diluted 1:40 for 30min. Then, tissue was again washed 2x 5 minutes using 1x Wash Buffer A with gentle shaking. Samples for Fluorescent-PLA were incubated for 1.5h with 1x Duolink^®^ Amplification Buffer (Green) containing the polymerase diluted 1:80. After 2x washes for 10 min with 1x Wash Buffer B and gentle shaking, sections were coverslipped with mounting medium (Prolong Gold) and imaged using a Zeiss confocal microscope with an 880 detector and a 63x oil lens. Samples for BF-PLA were incubated for 1.5 hours with 1x Duolink^®^ Amplification Buffer (Brightfield) containing the polymerase diluted 1:80. After amplification, tissue was washed 2x 2min with 1x Wash Buffer A using gentle shaking. Samples were further incubated in 1x Detection Brightfield for 1 hour at room temperature. After 2 washes for 2 min with Wash Buffer A and gentle shaking, sections were incubated in Developing Solution containing Substrate Reagent A (1:70), Reagent B (1:100), Reagent C (1:100), and Reagent D (1:50) for 15min at room temperature. Following 2 washes for 2 minutes in Wash Buffer A, tissue was dehydrated using increasing alcohol solutions and Xylene, and coverslipped using DPX mounting media. Sections were imaged using a Keyence All-In-One microscope with a 10x or 63x oil lens. Z-stacks were merged using the standard full focus option.

## Acknowledgements

This research was supported in part by the Intramural Research Program of the NIH, National Institute on Aging, by the Michael J. Fox Foundation Grant 16376, by the NIMH IRP Rodent Behavioral Core (ZIC MH002952 and MH002952 to Yogita Chudasama), by the NICHD Microscopy & Imaging Core, the NINDS viral production core facility and the NHLBI Electron Microscopy Core. The authors also want to thank Dr. Jillian Kluss for her support and input on the PLA protocol.

## Supplementary Figure 1

**Supplementary Figure 1:**
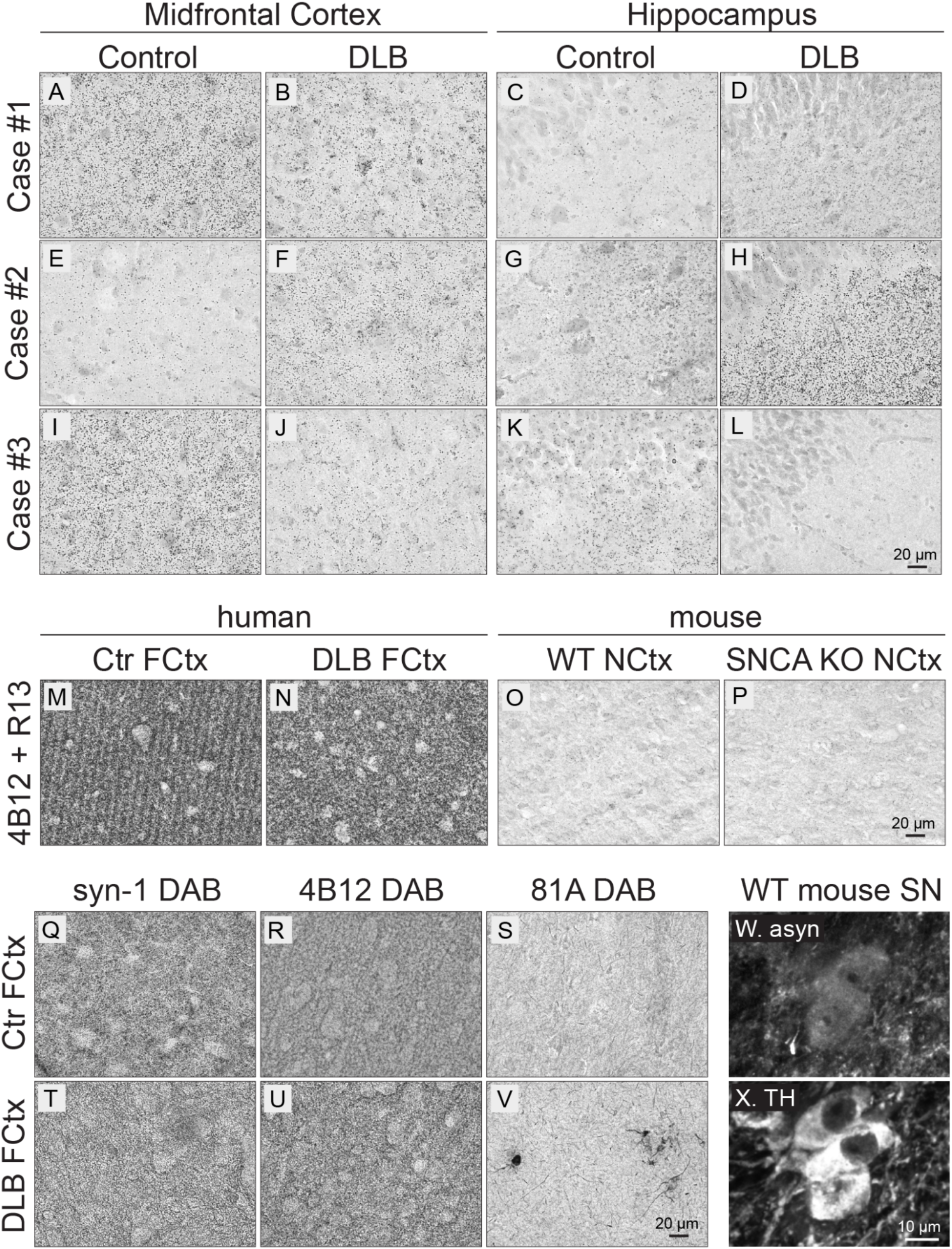
Verification of human cortical BF-PLA staining. Three additional human DLB cases (**B,F,J,D,H,L**) and three healthy controls (**A,E,I,C,G,K**) were stained using the BF-PLA (**A-L**). Sections from the Midfrontal Cortex (**A,B,E,F,I,J**) and Hippocampus (**C,D,G,H,K,L**) were chosen. BF-PLA was also performed on frontal cortical human (**M,N**) and neocortical mouse sections (negative control; **O,P**) using the human-specific total asyn antibody 4B12 and the previously used pS129 asyn antibody R13. Traditional single antibody DAB staining was used to evaluate the ability of syn-1 (**Q,T**), 4B12 (**R,U**) and 81A (S,V) to detect LB in frontal cortical tissue of healthy control (**Q-S**) and DLB cases (**T-V**). Mouse nigral sections were stained for asyn (syn-1; **W**) and Tyrosine Hydroxylase (TH; **X**) to identify dopaminergic neurons. All images are taken at 63x using either the Keyence All-In-One microscope (A-V) or the Zeiss LSM 880 confocal microscope (**W,X**).

